# Spatial Models of Pattern Formation During Phagocytosis

**DOI:** 10.1101/2022.04.11.487979

**Authors:** J. Cody Herron, Shiqiong Hu, Bei Liu, Takashi Watanabe, Klaus M. Hahn, Timothy C. Elston

**Affiliations:** Curriculum in Bioinformatics and Computational Biology, University of North Carolina at Chapel Hill, Chapel Hill, NC, USA; Computational Medicine Program, University of North Carolina at Chapel Hill, Chapel Hill, NC, USA; Department of Pharmacology, School of Medicine, University of North Carolina at Chapel Hill, Chapel Hill, NC, USA; Cancer Center Division of Gene Regulation, Fujita Health University, Toyoake, Japan

## Abstract

Phagocytosis, the biological process in which cells ingest large particles such as bacteria, is a key component of the innate immune response. Fcγ receptor (FcγR)-mediated phagocytosis is initiated when these receptors are activated after binding immunoglobulin G (IgG). Receptor activation initiates a signaling cascade that leads to the formation of the phagocytic cup and culminates with ingestion of the foreign particle. In the experimental system termed “frustrated phagocytosis”, cells attempt to internalize micropatterned disks of IgG. Cells that engage in frustrated phagocytosis form “rosettes” of actin-enriched structures called podosomes around the IgG disk. The mechanism that generates the rosette pattern is unknown. We present data that supports the involvement of Cdc42, a member of the Rho family of GTPases, in pattern formation. Cdc42 acts downstream of receptor activation, upstream of actin polymerization, and is known to play a role in polarity establishment. Reaction-diffusion models for GTPase spatiotemporal dynamics exist. We demonstrate how the addition of negative feedback and minor changes to these models can generate the experimentally observed rosette pattern of podosomes. We show that this pattern formation can occur through two general mechanisms. In the first mechanism, an intermediate species forms a ring of high activity around the IgG disk, which then promotes rosette organization. The second mechanism does not require initial ring formation but relies on spatial gradients of intermediate chemical species that are selectively activated over the IgG patch. Finally, we analyze the models to suggest experiments to test their validity.

**Author Summary:** Phagocytosis, the process by which cells ingest foreign bodies, plays an important role in innate immunity. Phagocytosis is initiated when antibodies coating the surface of a foreign body are recognized by immune cells, such as macrophages. To study early events in phagocytosis, we used “frustrated phagocytosis”, an experimental system in which antibodies are micropatterned in disks on a cover slip. The cytoskeleton of cells attempting to phagocytose these disks organizes into “rosette” patterns around the disks. To investigate mechanisms that underlie rosette formation we turned to mathematical modeling based on reaction-diffusion equations. Building on existing models for polarity establishment, our analysis revealed two mechanisms for rosette formation. In the first scenario an initial ring of an intermediate signaling molecule forms around the disk, while in the second scenario rosette formation is driven by gradients of positive and negative pathway regulators that are activated over the disk. Finally, we analyze our models to suggest experiments for testing these mechanisms.

## Introduction

All cells must be able to respond to changes in their environment, and often the proper response requires cells to adopt a new morphology. For example, cell shape changes occur during migration, division, and phagocytosis. Typically, these changes are initiated when receptors on the cell surface are activated by an external cue [1]. Receptor activation initiates a signaling cascade that results in spatiotemporal regulation of the actin cytoskeleton. The Rho family of GTPases are a class of signaling molecules that play key roles in this process [2]–[6]. These proteins act as molecular switches. They are in an inactive state when bound with GDP and become active when GDP is exchanged for GTP. Once active, Rho GTPases interact with effector molecules including those that regulate the actin cytoskeleton. Due to the nonlinear nature of the signaling pathways that regulate GTPase activity, understanding the molecular mechanisms that generate cell shape changes has proven challenging [1]. Therefore, many recent studies have turned to mathematical modeling to explore mechanisms capable of generating complex molecular structures [7]–[11].

Here we focus on Fcγ Receptor (FcγR)-mediated phagocytosis because of its biological importance in the innate immune response [12], [13] and because phagocytosis provides an outstanding system for studying how Rho GTPases organize the cytoskeleton into well-defined structures. Phagocytosis is initiated by the binding of the antibody Immunoglobulin G (IgG) to FcγR. Upon FcγR clustering, receptor cross-linking leads to phosphorylation of activation motif domains, enabling downstream signaling [12]–[14]. To study the events that initiate phagocytosis under well-controlled conditions, IgG is micropatterned in small disks on a glass coverslip (Fig. 1A). Because the antibody is attached to the coverslip it cannot be internalized, and the experimental system is therefore referred to as “frustrated” phagocytosis [15].

**Figure 1.**
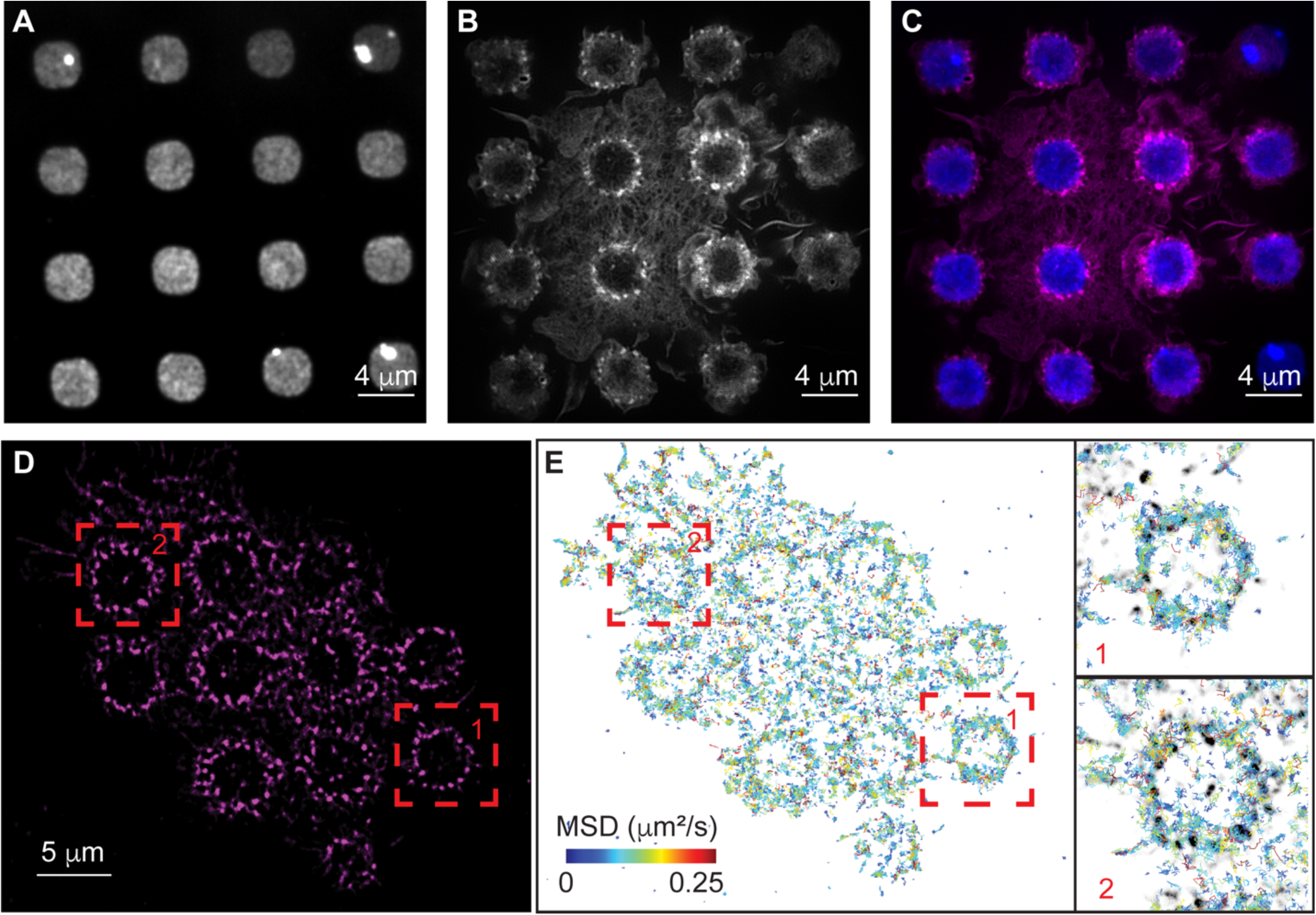
Actin and Cdc42 localization around disks of IgG. **A-C)** Rosettes of actin podosomes form around IgG disks. A micropattern of IgG disks of diameter 3.5 μm is shown in **A**. Actin is shown in **B**, in which puncta are podosomes. An overlay (IgG in blue, actin in magenta) is shown in **C**. **D, E)** Single particle tracking of Cdc42 during frustrated phagocytosis. Actin is shown in **D**. Single particle tracking of Cdc42 is shown in **E**, with tracks colored by their mean squared displacement. Inserts in **E** show two individual examples corresponding to the red boxes, with MSD colored as in **E** and actin shown in grayscale.

Following receptor activation, actin-enriched, adhesion-like structures termed podosomes [12], [13], [16] form in a circle around the IgG disk (Fig. 1B,C). Podosomes recruit many additional molecules and are thought to coordinate interactions between the actin cytoskeleton and the extracellular matrix [16]–[18]. They also form the leading edge of the phagocytic cup [19], [20]. The mechanisms responsible for podosome formation and patterning are not known. Therefore, we turned to mathematical modeling to establish sufficient conditions for pattern formation during frustrated phagocytosis.

Beginning with Turing’s seminal paper [21] and continuing with developments by Gierer and Meinhardt [22] and Meinhardt [23], reaction-diffusion models have been used to investigate pattern formation in biological systems. These models rely on positive feedback to amplify local fluctuations in signaling activity and some form of global inhibition to keep regions of high activity localized [22]. Another key requirement of these models is that at least one of the chemical species in the system diffuses at a different rate from the others [21], [22]. The hydrolysis cycle of GTPases satisfies the requirements for spontaneous polarization [7]–[10], [24], [25]. GTPases cycle between an active state when GTP-bound and an inactive state when GDP-bound. Their activation is catalyzed by guanine nucleotide exchange factors (GEFs), which promote the exchange of GDP to GTP. This exchange typically occurs at the cell membrane where diffusion is slow as compared to the cytosol [7], [8], [10], [25], [26]. When in the active state, some GTPases have been shown to recruit their own GEFs forming a positive feedback loop [24], [25], [27]–[30]. GTPase inactivation is accelerated by GTPase-activating proteins (GAPs) [3]–[6]. When inactive, GTPases are sequestered in the cytosol by guanine nucleotide dissociation inhibitors (GDIs) and diffuse rapidly [3]–[6].

There are now many reaction-diffusion models that describe how GTPases can generate cell polarity and patterning in various systems [8]–[11]. One of the best characterized cases is in yeast (*Saccharomyces cerevisiae*) budding, in which the GTPase Cdc42 generates a single, active site to determine the location of a bud site or mating projection. In yeast, autocatalysis is well-defined: active Cdc42 binds to the scaffold protein Bem1, which subsequently binds to the GEF Cdc24 that locally activates more GTPase [24], [25], [27], [28]. Another well characterized system is in single-cell wound healing, where Rho and Cdc42 form in distinct rings through the dual GAP-GEF Abr [31], [32]. Other examples include tip growth in pollen tubes and fungal hyphae [10] and cell motility [33], [34].

Here, we expanded upon existing reaction-diffusion models for GTPase activity to demonstrate how these systems can generate the “rosette” pattern of podosomes observed during frustrated phagocytosis. We explore the behavior of a recent model for GTPase activity that includes a negative feedback loop formed through the activation of a GAP [8]–[11]. Depending on the choice of parameter values, this model generates a range of patterns including spots, mazes, and inverse spots. We use the model to identify two distinct mechanisms for generating a rosette pattern. In the first scenario, an intermediate species forms a ring of activity that promotes the formation of active GTPase spots in the ring. Next, we use a parameterization approach involving an evolutionary algorithm followed by Markov chain Monte Carlo to evolve systems that do not rely on initial ring formation to generate the rosette pattern. A common theme that emerges from this analysis is that rosette formation requires the activation of both a positive and negative regulator of GTPase activity over the IgG disk. This creates spatial gradients of these regulators, which in turn are sufficient to drive the formation of the rosette pattern. Finally, we analyzed the behavior of the models to suggest experiments to test our proposed mechanisms.

## Results

### Experimental observations suggest Cdc42, but not myosin, is required for rosette patterning

Macrophages (RAW 264.7 cells) were observed during frustrated Fcγ receptor IIa (FcγR) mediated phagocytosis, where cells attempt to phagocytose fixed, micropatterned disks of immunoglobulin G (IgG). Actin, a major downstream effector during FcγR-mediated phagocytic signaling, formed in rings of small puncta, just outside of the IgG disks (Fig. 1A-C). These puncta were podosomes: actin-rich, adhesion-like structures observed during phagocytosis but more commonly known for their roles in motility and extracellular matrix interactions [16], [17]. This superstructural organization of podosomes in a circular arrangement has previously been termed a podosome “rosette” [35]–[37]. Due to the dynamic nature of phagocytosis, actomyosin contractility is known to play an integral role during the engulfment process [12], [13], [20], [38] and myosin II has been observed to localize to phagocytic podosomes and podosome rosettes [18], [20]. Therefore, we wondered whether actomyosin contractility was important for podosome rosette formation. To test this possibility, we treated cells with the Rho kinase inhibitor Y27632. Inhibition of Rho kinase during frustrated phagocytosis led to the complete disassembly of myosin II filaments, demonstrating that myosin II contractility was inhibited (Fig. S1). However, podosome rosettes still formed (Fig. S1), which suggested that the formation and maintenance of podosomes during phagocytosis is independent of actomyosin contractility and that a biochemical mechanism may underlie rosette formation.

Rho family GTPases, including Cdc42 are known to be activated during FcγR-mediated phagocytic signaling [2], [12], [13], [39], [40]. Cdc42 is a regulator of the actin cytoskeleton, so we next examined its localization during frustrated phagocytosis. Cdc42 was visualized during frustrated phagocytosis using single particle tracking (Fig. 1D,E). Cdc42 appeared to colocalize to the podosome rosette with individual tracks observed near podosomes (Fig. 1E).

Taken together these results suggest podosome rosette organization involves localized Cdc42 activity but does not require active myosin-mediated force generation. Cdc42 is known to play a role in cell polarization. Therefore, we decided to investigate if a similar mechanism might underlie formation of the podosome rosette.

### Forming coexistent clusters of active GTPase

The core components of mathematical models for polarity establishment include an inactive form of a GTPase that is cytosolic and diffuses rapidly, an active form that is membrane bound and diffuses slowly, and positive feedback through autoactivation [7]–[11]. Additionally, these models often assume that mass is conserved and, therefore, do not include protein synthesis and degradation. In their simplest form, these models typically form a single polarity site [7]–[10], [24]. Recent investigations have focused on establishing mechanisms that generate coexisting active sites. For example, Chiou et al. [8] demonstrated how local depletion increases the competition time between clusters so that coexistence is maintained over biologically relevant time scales. Jacobs *et al.* [10] found that either adding protein synthesis and degradation or adding negative feedback through a GTPase-activating protein (GAP) could limit the growth of active clusters of GTPase, thus enabling coexistence. Here, we focused on one of the GAP models that balances biological relevance with mathematical simplicity.

The Wave-Pinning GAP model (WPGAP, Fig. 2A, [10]) is described mathematically by the following set of reaction-diffusion equations:

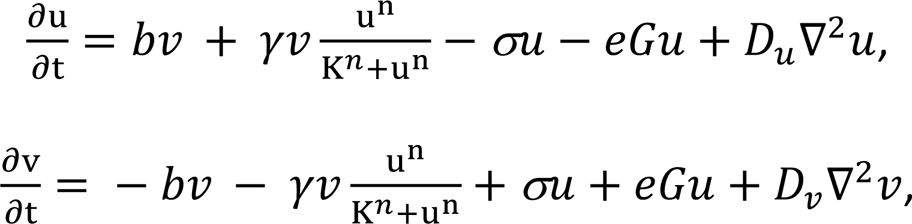

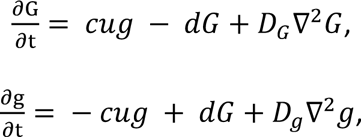

**Figure 2.**
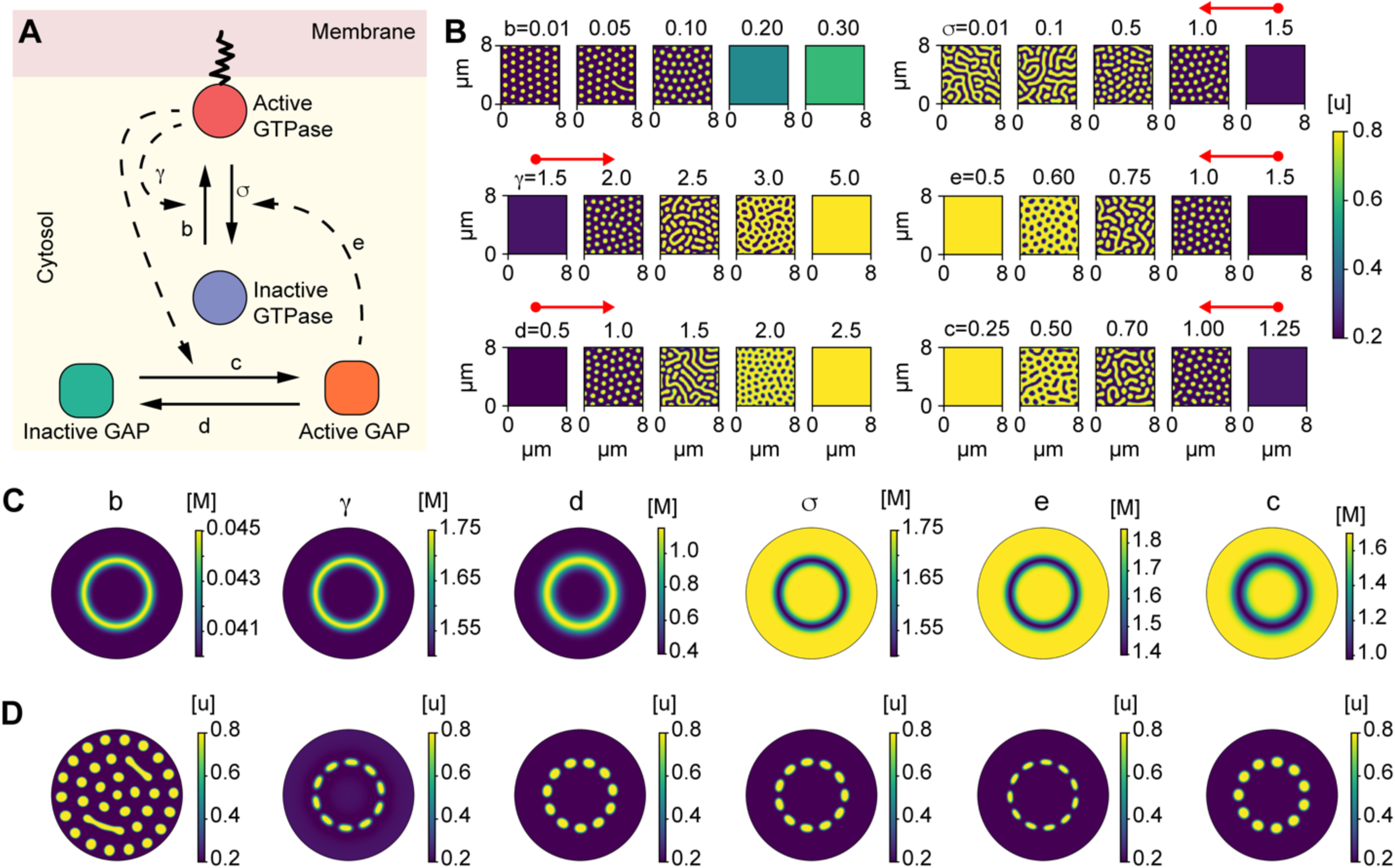
The WPGAP model generates rosettes of GTPase activity if a ring of an intermediate species is assumed. **A)** Schematic of the Wave-Pinning GTPase Activating Protein (WPGAP) model. **B)** WPGAP simulation results for individual parameter sweeps (Table 1). Most parameters show a transition from a low intensity homogenous regime to a spot patterning regime (red arrows). For details on concentration units, see Methods. **C)** Radial distributions for a modulating species M. The ring shown in these panels represents either a region of high (first 3 panels) or low (last 3 panels) [M]. **D**) Active GTPase concentrations using the distribution for [M] shown in **C** (above each panel, respectively).

**Table 1.** Baseline parameter values (See Methods for explanation on arbitrary units)

| Parameter | Description | Value | Ref. |
| --- | --- | --- | --- |
| $b$ | GTPase activation | $0.1 \text{ s}^{-1}$ | [10] |
| $\gamma$ | GTPase maximal self-positive feedback rate | $2 \text{ s}^{-1}$ | [10] |
| $K$ | Half-maximal response GTPase concentration | 1 a.u.* | [10]/This study |
| $n$ | Hill coefficient | 2 | [10] |
| $\sigma$ | GTPase inactivation | $1 \text{ s}^{-1}$ | [10] |
| $c$ | GAP activation | 1 a.u. | [10] |
| $d$ | GAP inactivation | $1 \text{ s}^{-1}$ | [10] |
| $e$ | GAP dependent GTPase inactivation | 1 a.u. | [10] |
| $D_u$ | Active GTPase diffusion | $0.004 \mu\text{m}^2\text{s}^{-1}$ | This study |
| $D_v$ | Inactive GTPase diffusion | $100D_u \mu\text{m}^2\text{s}^{-1}$ | [10]/ This study |
| $D_G$ | Active GAP diffusion | $100D_u \mu\text{m}^2\text{s}^{-1}$ | [10]/ This study |
| $D_g$ | Inactive GAP diffusion | $100D_u \mu\text{m}^2\text{s}^{-1}$ | [10]/ This study |
| $T$ | Total GTPase Concentration | 4.04 a.u. | [10] |
| $T_g$ | Total GAP Concentration | 10 a.u. | [10] |

where *u* is the concentration of active GTPase, *v* is the concentration of inactive GTPase, *G* is the concentration of active GAP, and *g* is the concentration of inactive GAP. The basal GTPase activation rate is *b*, the maximum self-positive feedback rate is γ, *K* is the concentration of active GTPase when the feedback is at the half-maximal response, the basal GTPase inactivation rate is *σ*, the GAP-mediated negative feedback rate is *e*, the GAP activation is *c*, and the GAP inactivation rate is *d*. The total mass of both species is conserved:

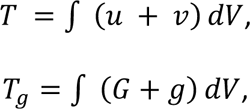

where the integrals are over the volume of the system. A requirement for polarization is that the membrane-bound active form of GTPase diffuses slowly in comparison to the cytosolic inactive form:

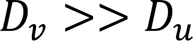

Adding negative feedback through GAP activation limits the growth of an individual cluster, allowing for coexistence of multiple clusters and other forms of patterning. Both the active and inactive forms of the GAP are treated as cytosolic species that diffuse rapidly compared to the membrane-bound active GTPase. Specifically, for the WPGAP model, Jacobs *et al.* [10] explored how changes in the total amount of GTPase (T) impacted bistability and the types of patterns that formed. They found the system could form spots (localized regions of high GTPase activity), mazes, and negative spots (localized regions of low GTPase activity). For our purposes, we are interested in coexisting spots, and thus we began with parameter values that placed the system in this regime (Table 1).

To gain insight into the model, we explored how spot size depends on the diffusion coefficients. By changing the diffusion coefficient of the active GTPase, *D_u_*, while holding the ratios for the other diffusion coefficients fixed, we found that increasing the diffusion coefficients produced an exponential increase in spot size (Fig. S2A,B). To demonstrate that changing the diffusion coefficients did not impact the patterns formed by the system, we also quantified the eccentricity for spots and saw no deviations from circularity (Fig. S2C). Radii of podosomes have been observed to be anywhere from ∼0.15 – 0.6 microns [18], [41]–[43]. Therefore, for our simulations we used *D_u_* = 0.004 *μm^2^ s^-1^*, which resulted in spots with radius 0.31 ± 0.02 µm (Fig. S2A-C).

Next, we explored how sweeping individual parameters changed pattern formation. When sweeping the GTPase activation rate *b*, lower values resulted in spots, but higher values resulted in a spatially homogenous steady state with an intermediate level of GTPase activity (Fig. 2B). This suggested that a finite basal GTPase activation rate *b* was not required or had to be quite small to facilitate patterning. Interestingly, each of the other parameter sweeps resulted in changes in the observed patterning types, including spots, mazes, and holes (Fig. 2B). For the GTPase inactivation rate *σ*, a high value resulted in a single low concentration throughout the domain, and decreasing this value led to spots, then mazes. However, minimizing this value did not cause the entire domain to be at a single, high steady state, due to negative feedback from GAPs. In contrast, the self-positive feedback rate *γ*, the GAP inactivation rate *c*, the GAP activation rate *d*, and the GAP-mediated negative feedback rate *e* were capable of all patterning types, from a single low state to spots, mazes, holes, and a single high state (Fig. 2B). Overall, these observations suggested that it would be possible to spatially modulate these parameters to go from the low, homogenous state to the spot forming state.

### A Two-Step Model for Rosette Formation

We next sought to determine if the WPGAP model could be modified to enable rosette formation. One possible explanation for how a rosette could form is if two distinct steps occur: 1) an initial ring of high or low concentration of some species (M) forms and 2) this species modulates a key parameter in the pattern forming, WPGAP model. To test this model, we first assumed that an initial ring forms. We discuss this assumption and potential mechanisms for ring formation later, but rings have been observed in other contexts, such as in wound healing, in which a chemical gradient and inhibition of a bistable GTPase by another resulted in two distinct rings of GTPase activity [31], [32]. For our initial investigations, we assumed that a modulator M affected a rate in the WPGAP model through the functional form:

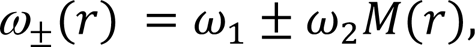

where *ω_1_* is the basal rate and *ω_2_* characterizes the effect of M on *ω* _±_. M(r) was modeled as a Gaussian-shaped function centered at r = 2 μm with variable variance. This form of *ω* _±_ allowed us to tune model parameters so that spot formation was only promoted within the ring.

For parameters that increase GTPase activity (GTPase activation *b*, GAP inactivation *d*, and the maximum self-positive feedback rate *γ*), the WPGAP model was coupled to a ring of high M concentration *ω*_+_ (Fig. 2C,D, three columns from the left). For parameters that decrease GTPase activity (GAP activation *c*, GTPase inactivation *σ*, and GAP-mediated GTPase inactivation *e*), the WPGAP model was coupled to an inverted ring of M, *ω*_-_ (Fig. 2C,D, three columns from the right). For each model parameter, *ω_1_* and *ω_2_* were varied to determine if the system could generate rosette organization. As an initial guess, the parameter values were chosen based on the results from the parameter sweeps (Fig. 2B).

As expected from the parameter sweep results, modulating the basal GTPase activation rate *b* did not appear sufficient to form a rosette pattern, because this produced spot formation throughout the entire domain. However, modulating the positive feedback rate *γ*, the GTPase inactivation rate *σ*, and the GAP-mediated negative feedback rate *e* all resulted in a rosette forming (Fig. 2C,D). Interestingly, when we modulated the rates for GTPase inactivation and the GAP-mediated negative feedback, we found that the rates required to form rosettes were higher than expected (Fig. 2C,D). For example, to form a rosette, the rate required for the GTPase inactivation σ within the ring was around 1.5, which resulted in no patterning when used as the global rate in the isolated WPGAP model (Fig. 2B-D).

### Gradient Establishment by a Simple Reaction-Diffusion Model

The analysis presented above demonstrated that the rosette pattern can form following the establishment of a ring of activity. Therefore, we next wanted to determine if rosette formation could occur in the absence of such an initial ring. To investigate this scenario, we first considered the following simple reaction-diffusion model:

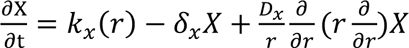

where X is a species that is activated with a rate *k_x_* that depends on the distance from the IgG patch. We assume that X is inactivated at a rate *δ_x_* and diffuses at a rate *D_x_* (Fig. 3A). Note that if *k_x_* is independent of r, then at steady state X(r) = *k_x_*/*δ_x_*. To model the IgG disk, we treat *k_x_(r)* as a step-function:

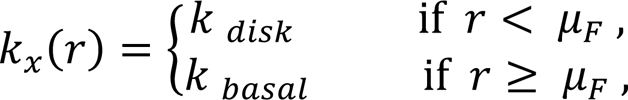

**Figure 3.**
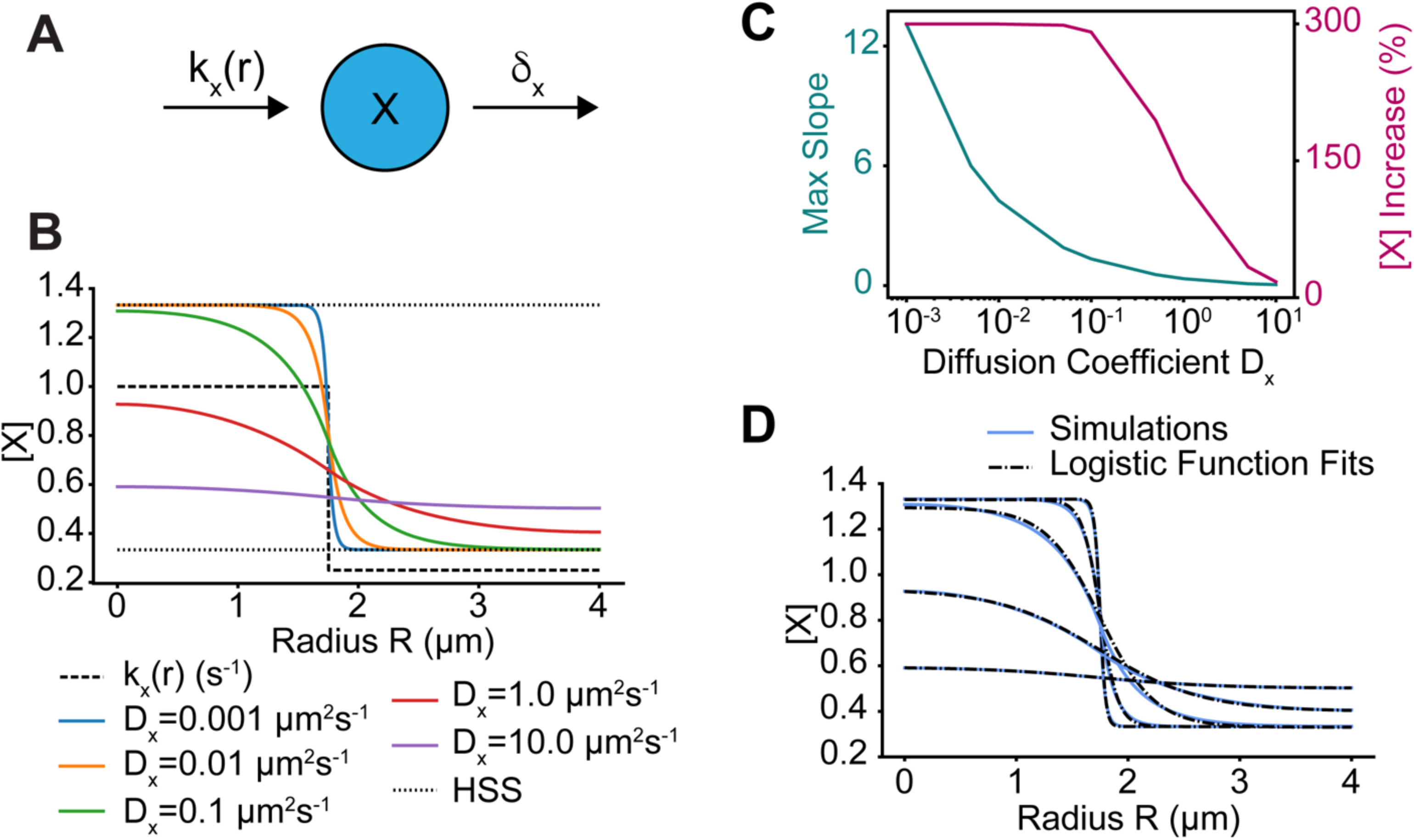
A simple reaction-diffusion model generates gradients of activity. **A)** Schematic of the simple reaction-diffusion model in which a species X is activated by a rate *k_x_(r)*, which depends on the local IgG concentration, and is deactivated at a constant rate *δ_x_*. **B**) Simulations of the model for various diffusion rates *D_x_*. The spatial profile of *k_x_(r)* is shown as the dashed line and *δ_x_* = 0.75 s^-1^. The homogenous steady states values of X when k = *k_x_(0)* and k = *k_x_(r_max_)* are shown as well (dotted lines). **C)** Maximum slope of X(r) and percent increase of X(0) over X(r_max_) as a function of *D_x_*. **D)** Blue curves are same as in **B** and dashed lines are best fits of these curves to the logistic function.

where *k_disk_* is the activation rate over a disk of radius *μ_F_* and *k_basal_* is the activation rate everywhere else. Thus, when *k_disk_* > *k_basal_*, this simple model describes a species whose activity is increased over the disk. For fixed values of *k_disk_*, *k_basal_*, and *δ_x_*, this system was simulated for varying diffusion coefficients *D_x_* (Fig. 3B). The range of diffusion coefficients was chosen to be consistent with reported cellular diffusion rates [26], from the slowest membrane-bound rate (10^-3^ μm^2^s^-1^) to the fastest cytosolic rate (10.0 μm^2^s^-1^). For a domain within r = 4 μm, changing the diffusivity of X resulted in changes in both the gradient steepness and the difference between the maximum and minimum values of X (Fig. 3C). For small values of *D_x_*, the distribution of X was switch-like, and X approached the expected steady state, *k_x_(r)*/*δ_x_*. However, for larger values of *D_x_*, the gradient in X was shallower, and deviated significantly from *k_x_(r)*/*δ_x_* with a lower total amplitude (Fig. 3B,C). We found that our simulation results could be well approximated using a logistic function with the form:

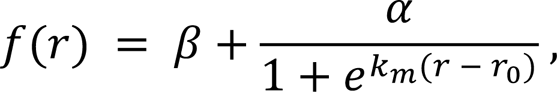

where *β* and *α+β* are the minimum and maximum values of f(r), respectively, *k_m_* is the logistic decay rate, and f(*r_0_*) = *β*+*α*/2 (Fig. 3D). The function f(r) was fit to distributions of X using simple optimization techniques (see Methods).

### Rosette Formation Through Gradients

To determine if rosette formation is possible in the WPGAP model without the formation of an initial ring around the IgG disk, we used the concentration profiles found above for the simple diffusion model to emulate activation over the disk. Thus, this system was the same as the WPGAP model, but the self-positive feedback rate *γ* and the GAP activation rate *c* now treated as spatially non-constant rates with profiles given by f(r).

Unlike the two-step model, it was difficult to empirically determine parameter values that form a rosette. Thus, we used a two-step approach to search parameter space. We first used an evolutionary algorithm (EA) [44] to perform a global search and subsequently performed a more local sampling of parameter space using a Delayed Rejection Adaptive Metropolis Markov chain Monte Carlo (DRAM-MCMC, see Methods) [45], [46].

To implement the two-step approach requires a score function that provides a quantitative measure for how close a simulated result is to the desired rosette pattern. For the desired pattern, a GTPase rosette formed by the two-step model was used (Fig. 2E). From this, we measured the radial average and radial standard deviation for the active GTPase *u* (Fig. 4A-C). For new simulations, we measured the radial average of *u* (Fig. 4A,C). We then divided the system into octants and measured the radial standard deviation within each octant (Fig. 4B,C). The difference between the means of the desired output and simulation result and the differences between the standard deviation of the desired output and standard deviations in each octant were calculated. These nine measurements (Fig. 4A-C) were then averaged to produce a single score. This score function was accurate, but flexible enough to allow for various numbers of spots and spot locations.

**Figure 4.**
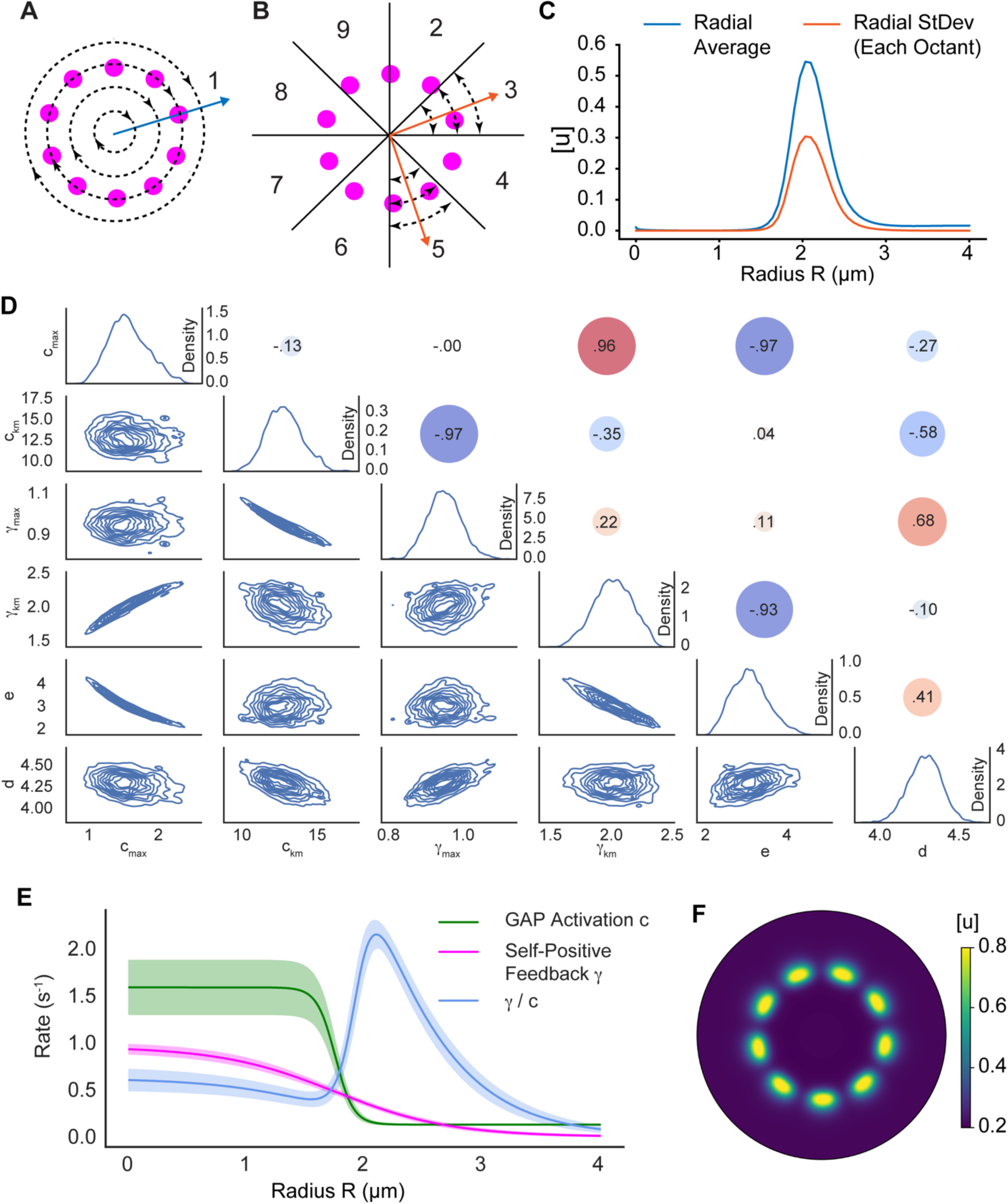
Model parameterization reveals mechanism enabling rosette formation. **A)** Schematic illustrating the radial averaging of GTPase activity used in the score function. **B)** Schematic illustrating the radial standard deviation of GTPase activity per octant used in the score function. This results in eight individual quantifications used in the score function. **C)** Radial profiles of the average and standard deviation of GTPase activity. These profiles were used in the score function and compared to the results from numerical simulations. **D)** Parameter distributions after performing DRAM-MCMC sampling. Individual parameter distributions are shown on the diagonal. Lower triangular plots show kernel density estimates for parameter pairs. Circles in the upper triangle represent the Spearman correlation coefficient between parameters. **E)** Radial profiles for the non-constant parameters from the parameter estimation. The positive to negative feedback ratio is also shown. The solid lines are the results for the mean parameter values from **D** (on the diagonal, Table 2) and the shaded regions indicate one standard deviation. **F)** Active GTPase concentration using the representative parameter set (Table 2). Simulation domain has a max radius of 4.0 μm.

**Table 2.**
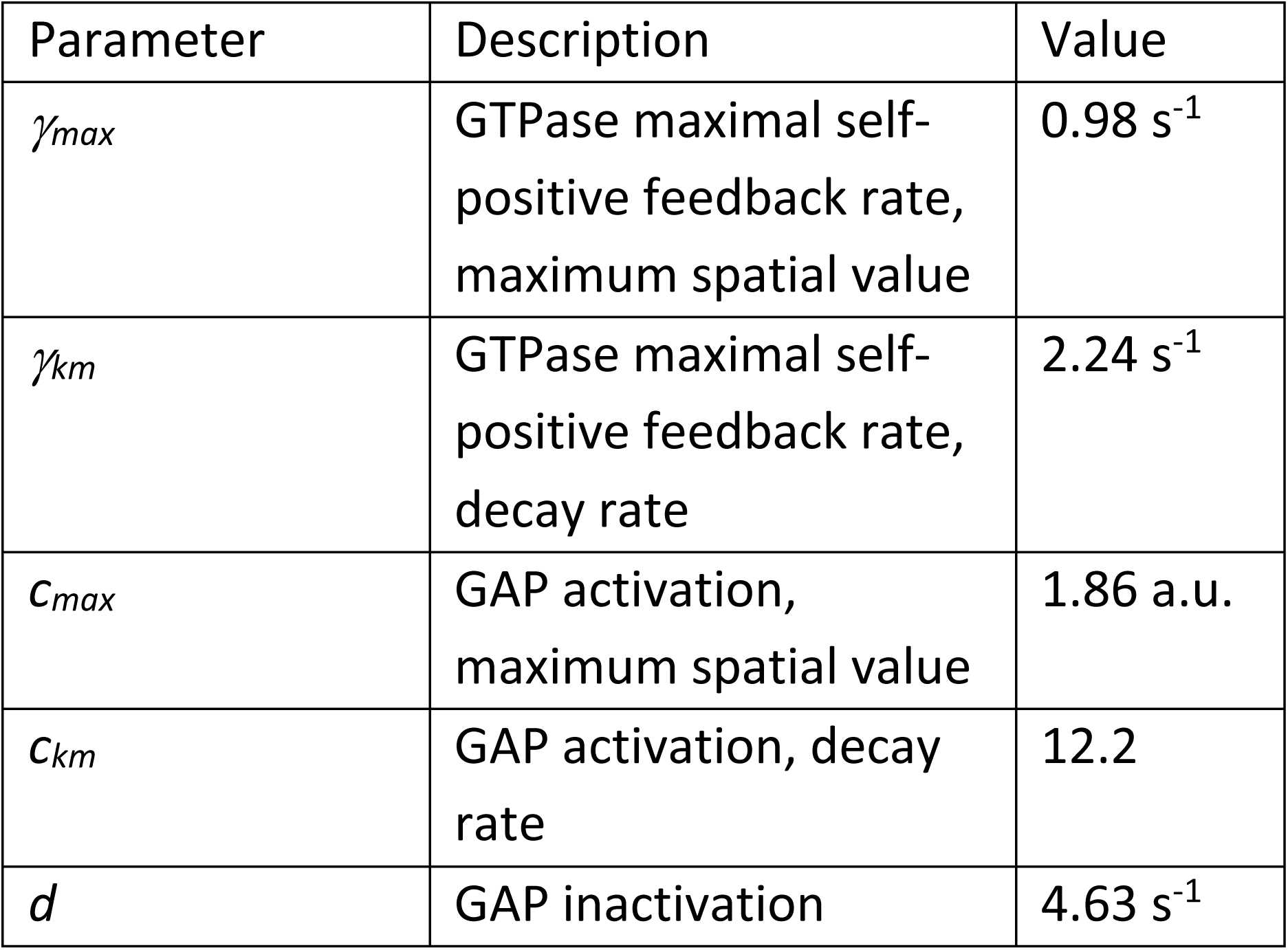
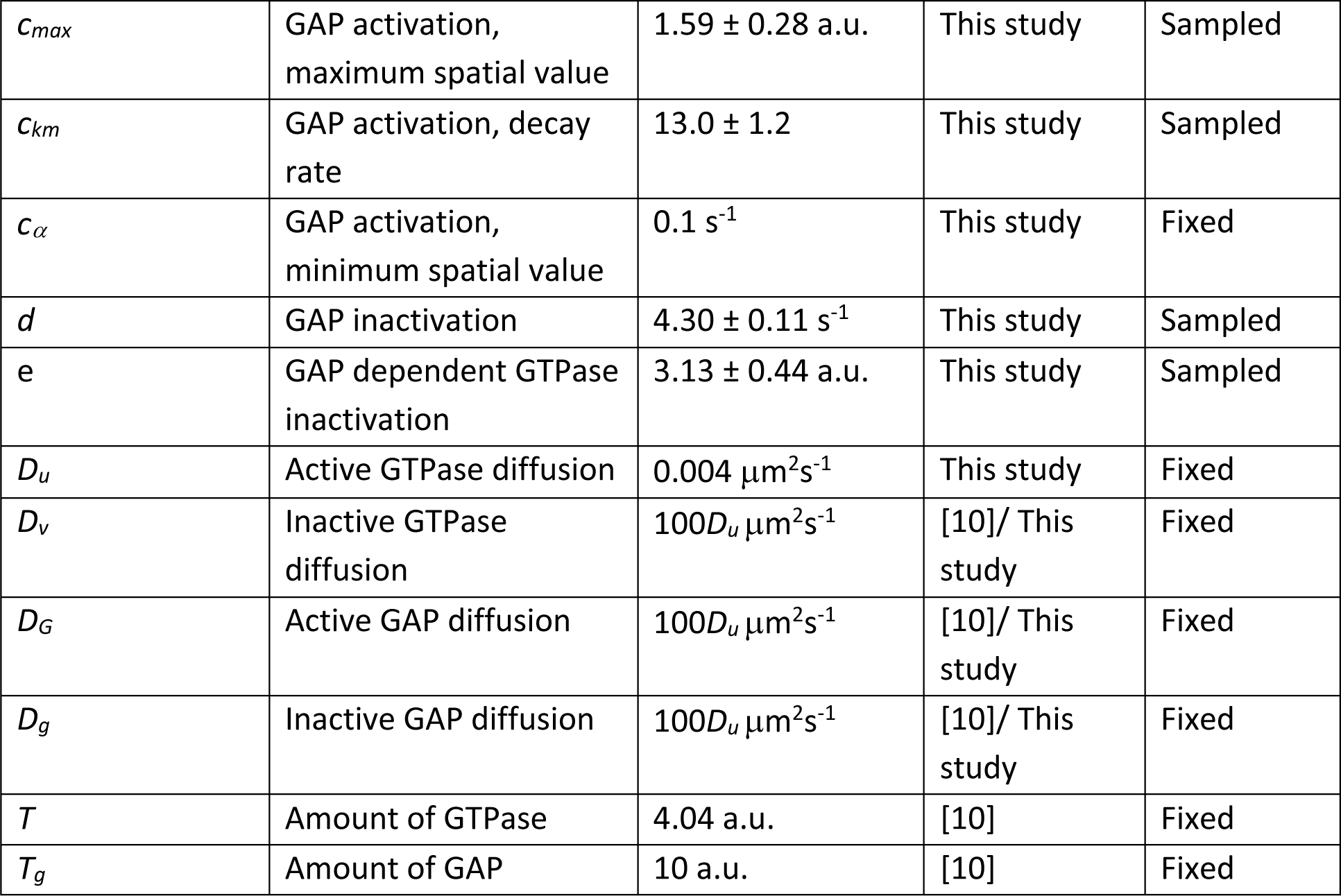
Mean parameter set after running the MCMC (See Methods for explanation on arbitrary units)

Simulations were initialized with a small amount of random noise and seeded with an initial concentration of active GTPase in the shape of a rosette (see Methods). We seeded simulations with a rosette to decrease the time for pattern formation, because parameter estimation requires a significant number of simulations. Also, from observations of initial parameterization attempts, the GTPase activation rate *b*, the GTPase inactivation rate *σ*, the minimum self-positive feedback rate *γ_α_*, and the minimum GAP activation rate *c_α_* were typically quite small and were thus fixed at 2e-4, 0.04, 5e-4, and 0.1 s^-1^, respectively. For 99 individual EA runs (100 individuals, 100 generations), most runs were able to discover parameters capable of rosette organization (top 80 appeared successful, Fig. S4A,B). The best parameter set found by the EAs was then used to initialize DRAM-MCMC simulations. DRAM-MCMCs were simulated until they appeared to converge, with all but the final 5,000 iterations removed as a “burn-in” period (Fig. S4C, see Methods).

The parameter distributions generated by MCMC sampling appeared Gaussian (Fig. 4D, on the diagonal). We took the mean values of the individual parameter distributions as our representative parameter set (Fig. 4D,F, Fig. S4D & Table 2). To check how well the MCMC performed, we also simulated the worst scoring parameter set, and these parameters also resulted in rosette organization (Fig. S4D,E & Table S1).

Inspection of the spatially dependent rates revealed how the system was capable of rosette patterning (Fig. 4E). When the ratio between the positive to negative feedback (*γ(r)*/*c(r)*) is plotted as a function of *r* using the identified parameter sets, in all cases the ratio is maximized just beyond r = 2 μm, near where the spots formed (Fig. 4E,F). The GAP activation rate *c(r)* (Fig. 4G, green) is high relative to the self-positive feedback rate *γ(r)* (Fig. 4G, magenta) over the disk and away from the disk. However, *c* transitions more rapidly than *γ* between its elevated level over the disk and its basal level away from the disk (Fig. 4G). Thus, while the negative feedback dominates over the disk and away from it, there is a zone near the edge of the disk where positive feedback surpasses negative feedback. It is in this region that rosette formation occurs.

To gain further insight into the model’s behavior we looked for pairwise correlations between model parameters. Several parameters demonstrated strong correlations (Fig. 4D). There was a strong anti-correlation between *c_max_*, the maximum GAP activation rate, and *e*, the rate constant for GAP-mediated GTPase inactivation. This likely indicates a sensitivity of the model to the total amount of GAP activity. The other correlations were not as intuitively apparent, so to further explore parameter-dependent model behavior, we performed individual parameter sweeps using the representative parameter set (Table 2, Fig. 5A-F). Parameters typically moved from no patterning to rosette organization to ring formation (*c_km_*, *γ_max_*, and *d*, Fig. 5B,C,E), or vice versa (*c_max_*, *γ_km_*, and *e*, Fig. 5A,D,F). Thus, the anti-correlations between *c_km_* and *γ_max_* as well as *γ_km_* and *e* likely result from a balancing of the effects of produced by varying the individual parameters. However, the reason for the positive correlation between *c_max_* and *γ_km_* is not readily apparent but may result from the logistic function having a lower maximum value for shallower gradients (i.e., low decay rates, Fig. 3).

**Figure 5.**
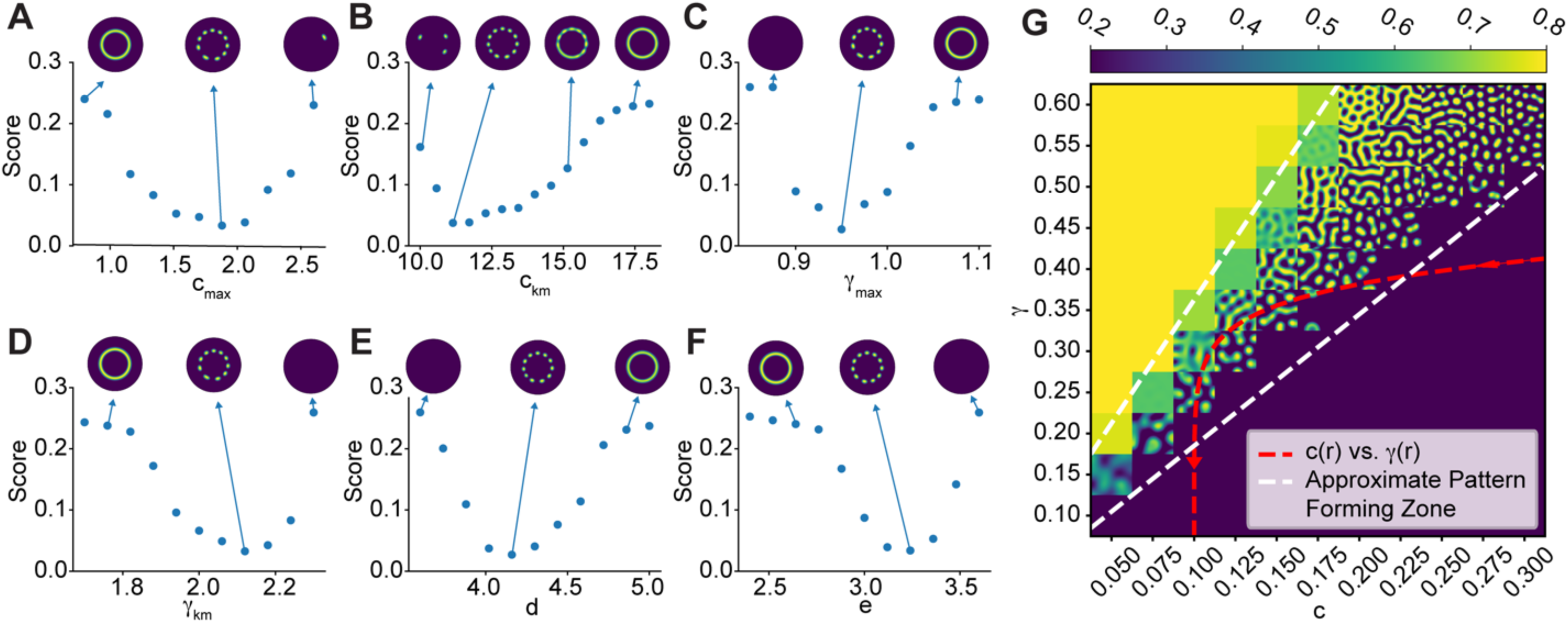
Effects of individual parameters on patterning. **A-F)** Plots of the score function versus model parameters (Table 2). Representative simulation results shown to illustrate the impact of varying parameter values on rosette formation. **G)** Two parameter sweep varying *c* and *γ*. Each grid cell shows the active GTPase concentration for an individual simulation when *c* and *γ* are fixed. The red dashed line is a plot of c(r) versus γ(r) from Fig. 4E & Table 2.

Finally, we performed a two parameter sweep for *γ* and *c*. We restricted the sweeps to the region of parameter space where spots formed (Fig. 4E,F). Individual simulations were performed using a constant value for *γ* and *c* (Fig. 5G). For high values of *γ* as compared to *c*, the model was in a high, homogenous regime (Fig. 5G, upper left). For high values of *c* as compared to *γ*, the model was in a low, homogenous regime (Fig. 5G, lower right). For intermediate values of *c* and *γ*, various types of patterning occurred, from spots to mazes to holes (Fig. 5G, bottom left to upper right). We next plotted *c(r*) vs. *γ(r)* within this region using the representative parameter set given in Table 2 (Fig. 5G, red curve). The curve is typically in the low, homogenous regime but passes through the patterning area, demonstrating why spots can form only within a certain spatial zone.

In the above simulations, the spatial profiles were prescribed using the logistic function. Therefore, we wanted to confirm that this mechanism would work for a coupled system, in which the modulating species were discretely modeled using the simple diffusion model with step-like activation over the IgG disk. The parameters in these two additional reaction-diffusion equations were optimized to fit the logistic functions for *γ(r)* and *c(r)* found above. Using the distributions for these two reaction diffusion-equations in the model did not lead to proper GTPase rosette formation, however there were differences between the logistic function fits and the simple reaction-diffusion equations, so this was not surprising. Therefore, we used the parameter values for these new equations to initialize another DRAM-MCMC. Because our goal was to simply demonstrate proof of principle, we only performed 22 short (1000 iterations) DRAM-MCMC runs, and we took the single best scoring parameter set. However, this was sufficient to demonstrate that a coupled model was able to generate a GTPase rosette (Fig. 6). In Fig. 6C and D, we plot the distributions of the species modulating the rates *c(r)* and *γ(r)*, respectively. Interestingly, if these profiles are used to modulate the intermediate species M in the two-step model, the system creates a ring of active M (as in Fig. 6E). Thus, the same mechanism where the positive feedback strength is lower, but transitions less rapidly than the negative feedback strength can also be used to create an initial ring which could then drive rosette formation. Finally, we note that with the right choice of parameter values, this model can also generate a ring, which could then be used to drive rosette formation (examples of this ring formation can be observed in Fig. 5A-F) as discussed above.

**Figure 6.**
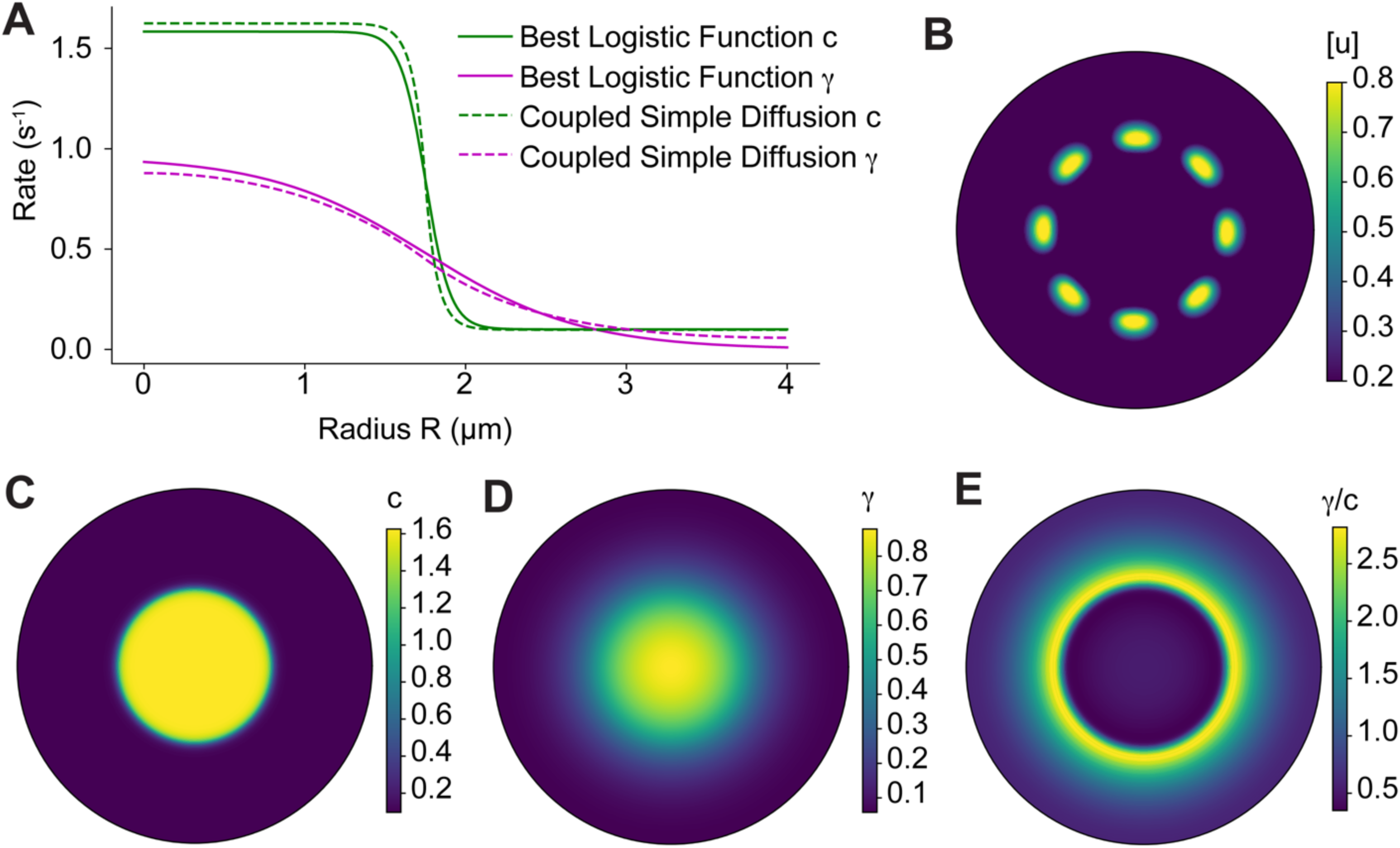
Rosette formation is possible when modulating species are explicitly included in the model. **A)** Radial distributions for *c* and *γ* from the coupled model (dashed lines) and the logistic function approximations using the representative parameter set (Table 2). **B)** Active GTPase concentration for the coupled model shown in **A**. **C)** Spatial concentration of the species modulating *c*. **D)** Spatial concentration of the species modulating *γ*. **E)** The positive to negative feedback ratio *γ/c* forms a ring.

### Experimentally Testable Predictions

Finally, we analyzed the model with the goal of motivating experimental investigations. First, we simulated the model using varying disk sizes (Fig. 7A,B). We observed a linear relationship between the disk radius and the number of spots, indicating that the distance between spots remains constant as the disk size increases (Fig. 7A). To compare our simulation results to experimental results, we counted the number of podosomes per site for IgG disks of radius 1.75 μm (8.1 ± 1.4, N = 29, Fig. S4A) and IgG disks of radius 5.0 μm (23.4 ± 2.4, N = 8, Fig. S4B). The observed number of spots qualitatively compared well with the numbers predicted by the model (Fig. 7B). For simulations using disk sizes of radius greater than 0.24 μm, 3 or more spots of active GTPase formed, while smaller disks formed 2 or fewer spots (Fig. 7A,B & S4C).

**Figure 7.**
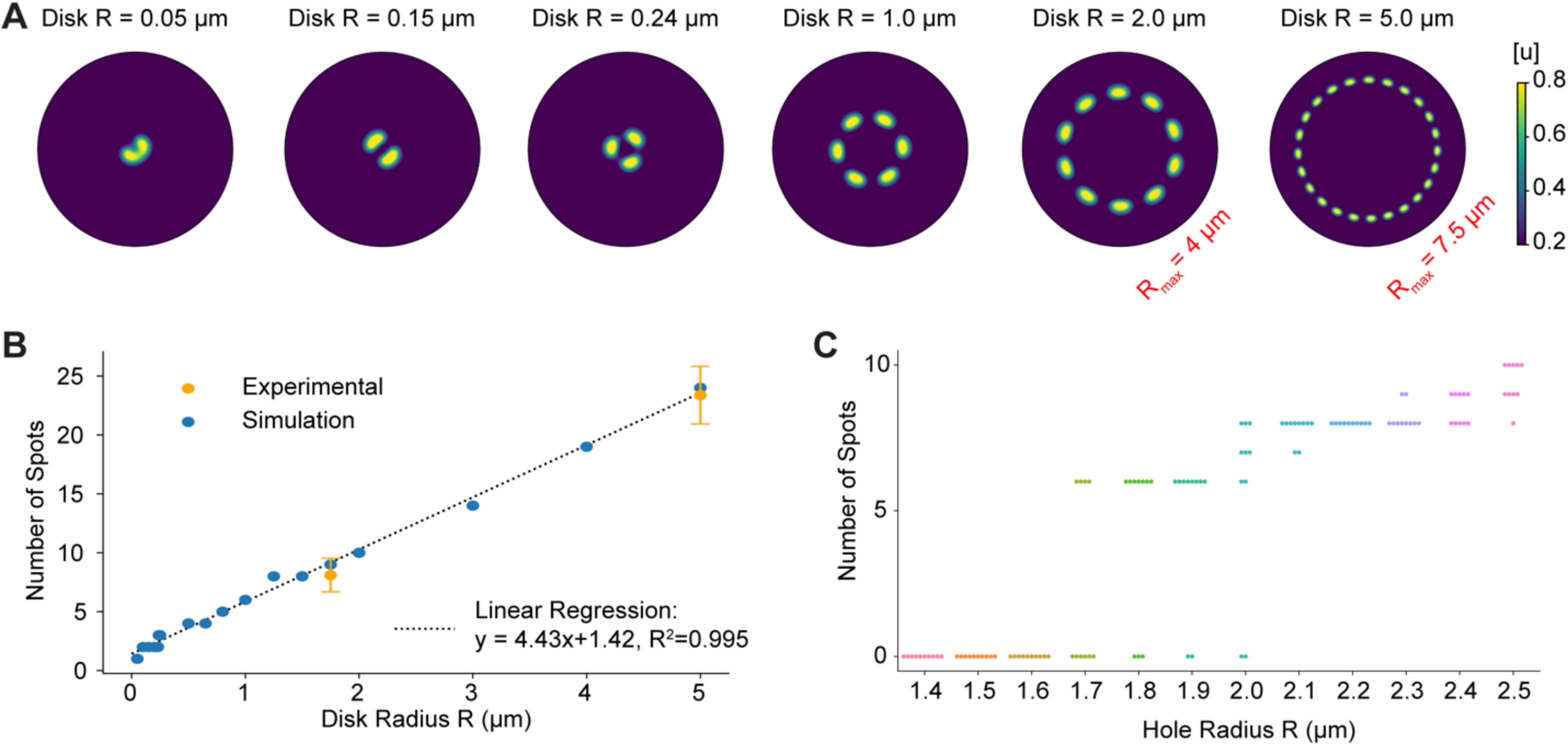
Effects of varying the size of IgG disks and holes. **A)** Simulation results for various disk using the representative parameter set (Table 2). The radius of the simulation domain is 4.0 μm except for the case with a disk size of R = 5.0 μm (far right), where the domain radius is 7.5 μm. **B)** The number of active GTPase spots linearly increases with disk sizes (blue circles). These results are consistent with experimental results (orange circles, whiskers denote one standard deviation). **C)** Number of active GTPase spots versus hole size. The system appears bistable for radii between 1.7 and 2.0 μm. In this region rosette formation depends on the initial conditions used in the simulations.

Similarly, we explored simulations with holes lacking IgG of various sizes (Table 2, negative *c_km_* and *γ_km_*, Fig. 7C & S4D). Rosettes always formed for a hole of radius 2.1 μm, and never formed for a hole of radius 1.6 μm. However, holes with radii between 1.7 and 2.0 μm appeared to be capable of forming rosettes, but rosettes sometimes failed to organize properly depending on the initial conditions, suggesting that the system is bistable within this regime (Fig. 7C).

## Discussion

Changes in cell morphology occur in many different physiological contexts, including cell migration, division, and differentiation. In eukaryotes, cell shape changes are driven by forces generated by the actin cytoskeleton. The Rho family of GTPases are primary regulators of the actin cytoskeleton. Therefore, understanding how these signaling molecules generate spatiotemporal patterns is fundamental to understanding cellular morphodynamics. An emerging theme in Rho GTPase signaling is that pattern formation occurs through a combination of positive autoregulation and differences in diffusivity between active and inactive GTPase states. It is now well understood how these elements can generate cell polarity (i.e., determining a cell front and back). However, what has been less clear is if this system can generate more complicated spatial patterns. Therefore, we turned to mathematical modeling to determine if this core polarity circuit can generate the rosette of podosomes observed during frustrated phagocytosis. Experimental evidence for such a biochemical mechanism comes from our observations that Cdc42 colocalizes to rosette structures and actomyosin contractility does not appear to be needed for rosette formation.

The starting point of our analysis was the WPGAP model that was previously shown to be capable of forming co-existing clusters of high GTPase activity [10]. The model consists of three main features: 1) active, membrane-bound GTPases diffuse slowly compared to cytosolic species, 2) active GTPases recruit other GTPase molecules from the cytosol to the membrane, and 3) a negative feedback loop is formed by activation of cytosolic GAPs. We used the model to investigate two potential mechanisms for rosette formation. In the first scenario, a ring of high or low concentration of a regulator of GTPase activity initially forms. Our analysis revealed that for rosette formation was possible if the species forming the ring regulated rates associated with positive feedback, GTPase inactivation, GAP-mediated negative feedback, GAP activation, or GAP inactivation. However, modulating the basal GTPase activation rate did not generate rosette formation, but instead spots of high GTPase activity formed throughout the entire domain.

We next used the model to demonstrate that rosette patterning could occur in the absence of initial ring formation. This scenario required the following conditions to be met: 1) the positive and negative feedback strengths increased over the IgG disk, 2) negative feedback dominated over positive feedback over the disk and far from the disk and 3) the negative feedback transitioned from its elevated level over the disk to its basal level more rapidly than that of the positive feedback. These three features generated a small region outside of the disk where the positive to negative feedback ratio is sufficiently high to enable spots of active GTPase. This scenario occurs if the positive regulator of GTPase activity diffuses rapidly, whereas the negative regulator of GTPase activity diffuses slowly in comparison. Furthermore, this same scenario could be used to generate a ring of high activity of species that positively regulates GTPase activity as required for the initial ring formation scenario described above.

Finally, we performed simulations using varying sizes of either IgG disks or holes. Experimental results for the number of spots formed using different IgG disk sizes were consistent with our simulation results. We also noticed that for simulations on disks of radius R = 0.24 μm, 3 distinct spots of GTPase activity were produced, whereas disks of smaller radii produced 1 or 2 sites. This result suggests a threshold for the minimum size of a particle that can be internalized via phagocytosis, if three or more podosomes are required to engulf a target. This observation is consistent with the reported value of 0.5 μm as the minimum size for phagocytic targets [47], [48]. Simulations on holes lacking IgG of different sizes revealed that holes of radii 1.6 μm or less do not form rosettes, while holes of 2.1 μm are capable of rosette formation. Interestingly, for holes between 1.7 μm and 2.0 μm rosette formation depended on initial conditions, suggesting the system is bistable in this regime.

Our analysis revealed how relatively minor additions to the Rho GTPase polarity circuit were sufficient to generate the rosette of podosomes observed during frustrated phagocytosis. It is likely that this same polarity circuit is also capable of generating more complex patterns when additional regulatory elements are added. Importantly, our results provide a computational framework for establishing sufficient conditions for more complex pattern formation, and therefore should be relevant to many different areas of cell biology. Here we focused on static cytoskeletal structures. However, the actin cytoskeleton is a dynamic system, and phagocytosis requires exact spatiotemporal control of cellular morphodynamics during engulfment. While this study demonstrated how initial cytoskeletal organization could occur, in future studies it will be important to also consider both the time-dependent and three-dimensional activity of Rho GTPase signaling during this process.

## Methods

### Cell Culture and Transfection

RAW 264.7 macrophages were obtained from the ATCC and maintained in culture medium: RPMI 1640 medium GlutaMAX Supplement (ThermoFisher Scientific, 61870127) containing 10% heat-inactivated FBS (HI-FBS, GEMINI Bio, 100-106) in a 5% CO_2_ humidified incubator at 37°C. To detach RAW 264.7 cells from the Falcon tissue culture dish (Fisher Scientific, 08-772E), the cells were treated with Accutase (ThermoFisher Scientific, A1110501) at 37°C for 5 min before gentle scraping (CytoOne, CC7600-0220). The plasmids FTractin-tdTomato and myosin regulatory light chain (MRLC)-EGFP were described previously [49], [50]. RAW 264.7 cells were electroporated with the Neon Transfection System (ThermoFisher Scientific) following the manufacturer’s protocol. In brief, 5×10^6^ cells were electroporated with 1 μg plasmid in R buffer at a setting of 1680 V, 20 ms, and 1 pulse using 10 microliter Neon pipette tip. The cells were transferred into a well of 12-well plate, with each well containing 1 ml of culture medium. After 12 hours of incubation, the transfected macrophages were ready for the frustrated phagocytosis experiments.

Bone marrow cells were isolated from 6 to 12 weeks C57BL/6 mice and differentiated into macrophages for 5-7 days in RPMI 1640 medium containing 10% heat inactivated FBS and 10% M-CSF (L929 conditioned medium) described elsewhere [51], [52]. These macrophages were detached from the flask using Accutase and gentle scraping.

### Microcontact printing

The IgG patterns on glass coverslips were made using the microcontact printing of Polydimethylsiloxane (PDMS) as previously described [53]. The silicon master with an array of 3.5 μm holes spaced 8 μm apart or 10 μm holes spaced 20 μm apart was made using photoresist lithography, and PDMS stamping on glass coverslips was carried out as described previously [43].

### Inhibition treatment and immunofluorescence staining

To inhibit actomyosin contractility and disassemble myosin II filaments, RAW 264.7 macrophages were plated on patterned IgG coverslips in Ham’s F12 medium (Caisson Labs, UT) supplemented with 2% HI-FBS and 20 μM Rho kinase inhibitor Y-27632 (Hello Bio, HB2297) for 25 min of inhibition during frustrated phagocytosis. For frustrated phagocytosis against 10 μm IgG spots, bone marrow-derived macrophages were plated on patterned IgG in the above medium without inhibitor and incubated at 37°C for 15 min before staining.

The cells were fixed with 4% paraformaldehyde at 37 °C for 10 - 15 min and permeabilized using 0.1% Triton-X-100 (Sigma-Aldrich) in PBS for 5 min. Cells were then thoroughly washed with PBS and fixative quenched with 0.1 M glycine for 20 min followed by incubation with 2% BSA fraction V (Thermo, 15260037) in PBS for 30 min. Actin was stained with Alexa-Fluor 568 phalloidin (dilution 1:500, ThermoFisher Scientific A12380) diluted in 2% BSA in PBS at room temperature for 20 min followed by one wash with 1xPBS/0.05% Tween for 10 min, and two washes with 1x PBS for 15 min.

### Imaging of podosome structures during frustrated phagocytosis

Total internal reflection fluorescence structured illumination microscopy (TIRF-SIM) was used to image podosomes in F-tractin-tdTomato transfected live RAW 264.7 macrophages. Fluorescence emission was recorded using an sCMOS camera (Hamamatsu, Orca Flash 4.0 v2 sCMOS). Lasers with wavelengths 560 and 647 nm and an Olympus UApo N 100x oil NA 1.49 objective were used, and fluorescence emission was recorded using an sCMOS camera (Hamamatsu, Orca Flash 4.0 v2 sCMOS). A Nikon SIM microscope was used to image podosomes in fixed RAW 264.7 macrophages after Y-27632 inhibition, using 488 and 561 nm lasers. A 100x oil immersion objective (1.49 NA, Nikon CFI Apochromat TIRF 100x) and EMCCD camera (Andor DU-897) were used. To image podosomes in bone marrow-derived macrophages, a Zeiss confocal microscope LSM880 built around AxioObserver 7 with a 63x 1.4 NA oil objective (Zeiss) was used.

### Single Particle Tracking

Single particle tracking was performed using a home-built total internal reflection microscope based on an Olympus IX81. The microscope was equipped with four solid state lasers (Coherent OBIS 405 nm, 488 nm, 561 nm, and 647 nm), a 100X TIRF objective (Olympus, UPLAPO100XOHR) and an sCMOS camera (Photometrics Prime 95B) for fluorescence collection. Raw cells were co-transfected with mScarlet-F-tractin (Excitation, 561 nm; Emission, Semrock, FF01-600/52) and Cdc42-HaloTag [54]. Cells were incubated with 100 pM dye JF646-Halo (Emission, Semrock, FF01-698/70) for 30 minutes and washed with culture medium three times before imaging. Super-resolved F-tractin images were acquired at 100 Hz for 5 seconds and subjected to Super-Resolution Radial Fluctuations analysis [55]. For single particle tracking of Cdc42, we streamed for 40 seconds at 50 Hz (2000 frames).

Single molecule diffusion analysis was done as before [56]. Briefly, individual molecules were identified by a wavelet decomposition based approach [57] and precise centroids were obtained by fitting with a 2D Gaussian function. Single molecule trajectories were built through a well-established linking algorithm [58] and the mean-square-displacement was then calculated [59], [60] to color encode the tracks.

### Numerical Simulations

Ordinary differential equation (ODE) simulations were performed by using the Python package odeint from Scipy [61]. Reaction-diffusion equations were solved using the spectral differential equation solver Python package Dedalus [62]. For simulations using Cartesian coordinates, the system was spatially discretized using a Fourier basis in *x* and a Chebyshev basis in *y* with the recommended dealiasing factor of 1.5, as done before [10]. The system had periodic boundary conditions in *x* and Neumann (reflective) boundary conditions in *y*. Similarly, for simulations using polar coordinates, the system was spatially discretized using a Fourier basis in *ϕ*, and a Chebyshev basis in *r* (dealiasing factor of 1.5) with periodic boundary conditions in *ϕ*, and Neumann (no flux) boundary conditions in *r*. The typical grid size used for simulations was 256 x 128 (*ϕ*, r respectively) which was informed by mesh grid refinement (i.e., larger grid sizes resulted in the same outcome). However, a grid size of 64 x 64 was used for parameterization steps to decrease simulation time. Simulations were typically performed using a time step dt = 0.1 s or 0.25 s. Reaction steps were solved using 4^th^ order Runge-Kutta, although 2^nd^ order Runge-Kutta was used for parameterization steps.

Homogeneous steady states were determined by running the ODE system (without diffusion) for *t* = 1000 s using odeint. For initial conditions of the reaction-diffusion equations, each species was set to its steady state value throughout the domain and subsequently noise was added by converting a small fraction of inactive species to the active form. The fraction of concentration converted was determined by each simulation but was typically generated by uniform sampling between 0 and 0.2*v_ss_* (where *v_ss_* is the steady state concentration for the inactive species). For seeded simulations, the same random noise (but between 0 and 0.1*v_ss_*) was converted and additionally the normalized seed (i.e., a rosette) was scaled by 0.1*v_ss_* and converted to active GTPase. For simulations with non-constant coefficients, the initial steady states were determined by using the basal values for the spatially-dependent rates.

To fit the logistic equation to the simple reaction-diffusion profiles, we used the Python package minimize from Scipy [61].

### Concentration Units

Because we currently do not know the average number of Cdc42 molecules associated with an individual podosome, we did not assign specific units to the total concentrations of Cdc42 and GAP. Thus, we used the unitless values from the non-dimensionalized version of the model described by Jacobs *et al.* [10], and labeled these as arbitrary units (a.u.). Note that once an estimate for the number of Cdc42 molecules per podosome is known, the total concentrations can be scaled appropriately to produce this number without changing any of our results. That is, scaling the concentrations for total Cdc42 and GAP by a factor χ and scaling the Hill constant K by χ and the second order rate constants *e* and *c* by χ^-1^ leaves the solutions to the reaction-diffusion equations scaled, but otherwise unchanged. For example, if χ = 100 then there would result in an average concentration of 404 Cdc42 molecules per μm^2^. Using our value for the radius of a podosome (0.31 μm), the area of a podosome is approximately 0.30 μm^2^. This would result in approximately 120 Cdc42 molecules per podosome. The units of concentration would then be molecules per μm^2^, and units of *K* would be the same. The second order rate constants *e* and *c* would have units of μm^2^(molecules*s)^-1^.

### Parametrization

Evolutionary algorithm (EA) simulations were performed using the Python package DEAP [44]. For EA hyperparameters we used a mutation rate of 0.3 and a crossover rate of 0.5. Markov chain Monte Carlo (MCMC) simulations were performed using the Python package Pymcmcstat [46]. For MCMC sampling, we used the Delayed Rejection Adaptive Metropolis (DRAM) algorithm [45], [46]. MCMC hyperparameters were set to S20 = 0.015 and N0 = 0.015, which resulted in chain acceptance rates between 29-40%. All but the last 5,000 steps for individual MCMC chains were discarded as a “burn-in” period. MCMC chains appeared to pass all convergence tests, including within chain variance (Geweke statistic *p* >> 0.05, [63]) and between chain variance (Gelman-Rubin diagnostic < 1.1, [64]).

Note that for the coupled model, where the simple reaction diffusion model was simulated in place of the logistic function, we used the same MCMC pipeline for sampling to discover a working coupled model. For proof of concept, we simply ran this pipeline for 1,000 steps and took the best scoring parameter set.

### Spot Size Determination

Simulations were performed as described above using cartesian coordinates (*t_final_* = 150 s). Each system was interpolated to a uniform grid with the same grid size (128 x 128). A mask was generated by thresholding at the mean of the maximum and minimum concentrations within the system. Using this mask, features were quantified using the Python package scikit-image [65]. The effective radius was defined as:

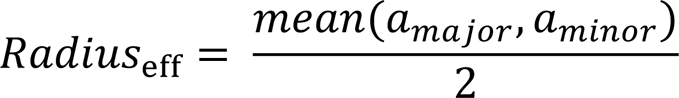

where *a_major_* and *a_minor_* were the major and minor axis lengths respectively. After an initial pass at fitting Radius*_eff_* based on log(*D_u_*), simulations were rerun using different grid sizes to ensure the number of spots counted for each simulation were similar (N∼100).

### Counting Podosomes Per Site

For experimental results, the number of podosomes per site were calculated using a pipeline we developed previously [43] (https://github.com/elstonlab/PodosomeImageAnalysis). In essence, this pipeline uses persistent homology, a type of topological data analysis, to identify significantly persistent features (connected components, holes) within images followed by post-processing.

For simulated results, a mask was created by thresholding at the average between the maximum and minimum intensity within a simulation. From this mask, the number of features was counted using the Python package ndimage.label from Scipy [61].

### Data Availability

The code and data used for this project is available on GitHub (https://github.com/elstonlab/PhagocytosisRosetteModel) and under the Zenodo archive: https://doi.org/10.5281/zenodo.6448430. Note that for counting podosomes, additional code is required from: https://github.com/elstonlab/PodosomeImageAnalysis.

## Acknowledgements

We thank Tony Perdue from the Department of Biology Microscopy Core at UNC for assistance with Nikon N-SIM microscopy, and Teng-Leong Chew, Aaron Taylor, Satya Khuon for assistance of TIRF-SIM and cell culture at Janelia Research Campus. We are grateful to Richard Superfine, Michael Falvo, and Timothy O’Brien for help with micropatterning and Ellen C. O’Shaughnessy, Wolfgang Bergmeier and Juan Song for primary macrophage culture. This work was supported by grants from the National Institute of General Medical Sciences (NIGMS) to KMH (R35GM122596) and to TCE (R35GM127145), as well as from the National Institute of Biomedical Imaging and Bioengineering to TCE (U01 EB018816). JCH received support from the NIGMS (5T32 GM067553).

## Contributions

SH, TW, and BL performed the experiments and imaging. BL performed the single particle tracking analysis. JCH performed the modeling and computational analyses. JCH and TCE wrote the manuscript with contributions from all authors. The work was directed by TCE and KMH.

---

**Supplemental Figure 1.**
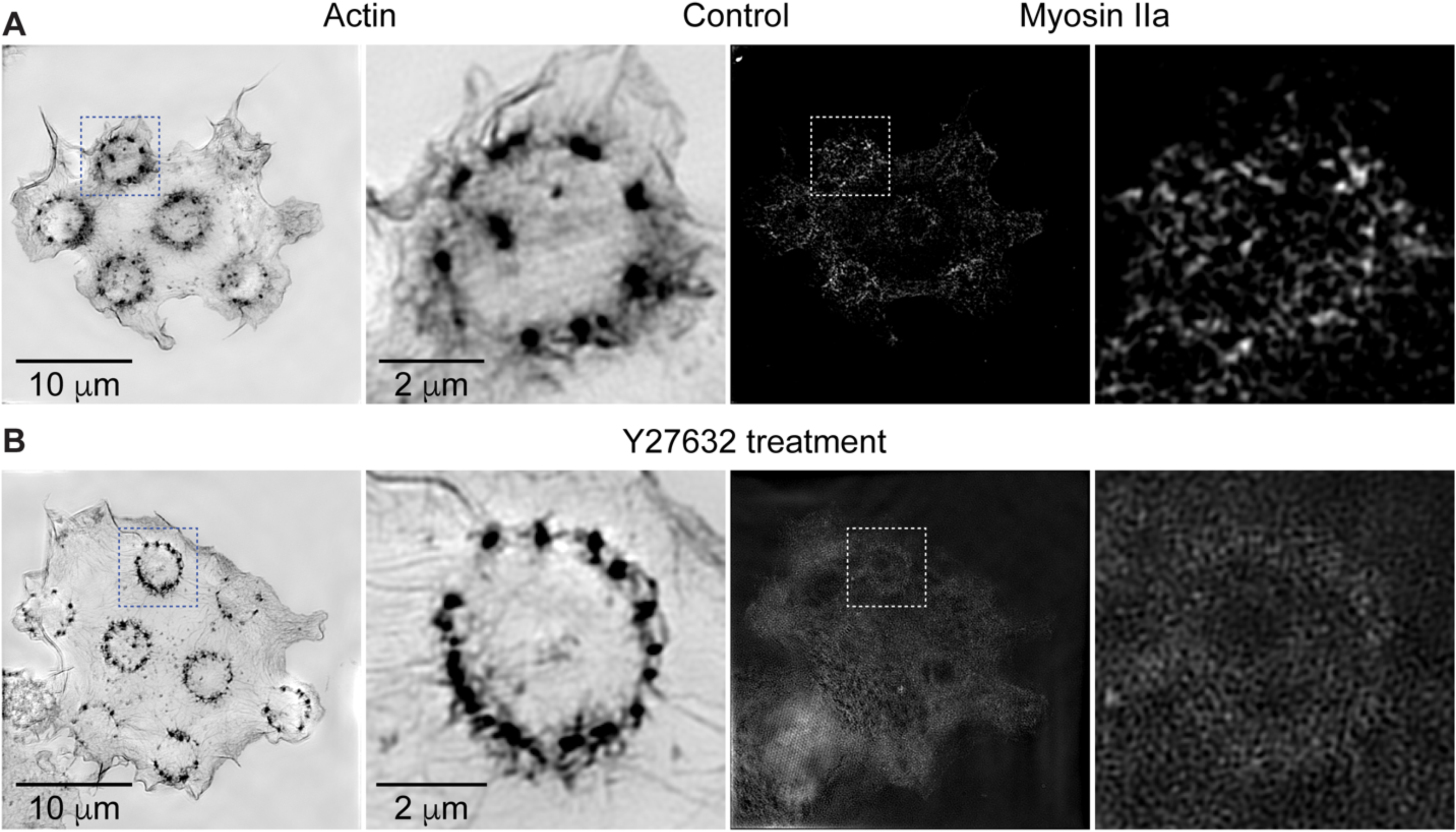
Actomyosin contractility does not appear to be necessary for rosette formation. **A**) Control RAW 264.7 macrophages were marked for actin (phalloidin staining) and myosin II (RLC-eGFP) during frustrated phagocytosis. **B**) RAW 264.7 macrophages were marked for actin (phalloidin staining) and myosin II (RLC-eGFP) when treated with 20mM Rho kinase inhibitor Y-27632 for 25 min during frustrated phagocytosis.

**Supplemental Figure 2.**
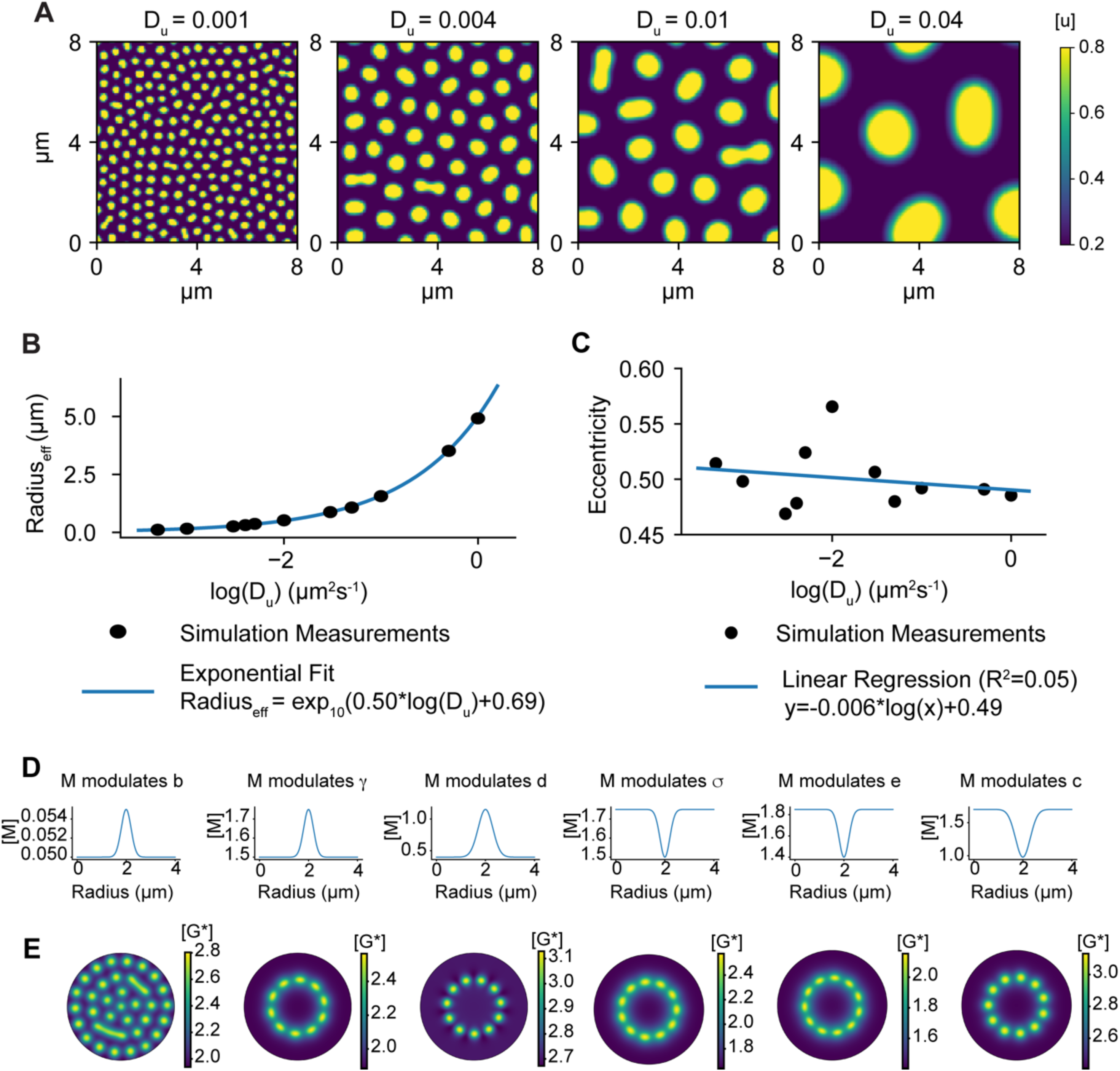
Diffusion rates determine the size of active GTPase spots. **A)** Simulations of the WPGAP model for variable diffusion rates. The diffusion coefficient for cytosolic species is taken to be 100*D_u_*. **B)** Relationship between the membrane diffusion coefficient and spot size. **C)** Relationship between the membrane diffusion coefficient and spot eccentricity. **D)** Radial distributions for the intermediate species M in Fig. 2E. **E)** Active GAP concentrations for the results shown in Fig. 2E.

**Supplementary Figure 3.**
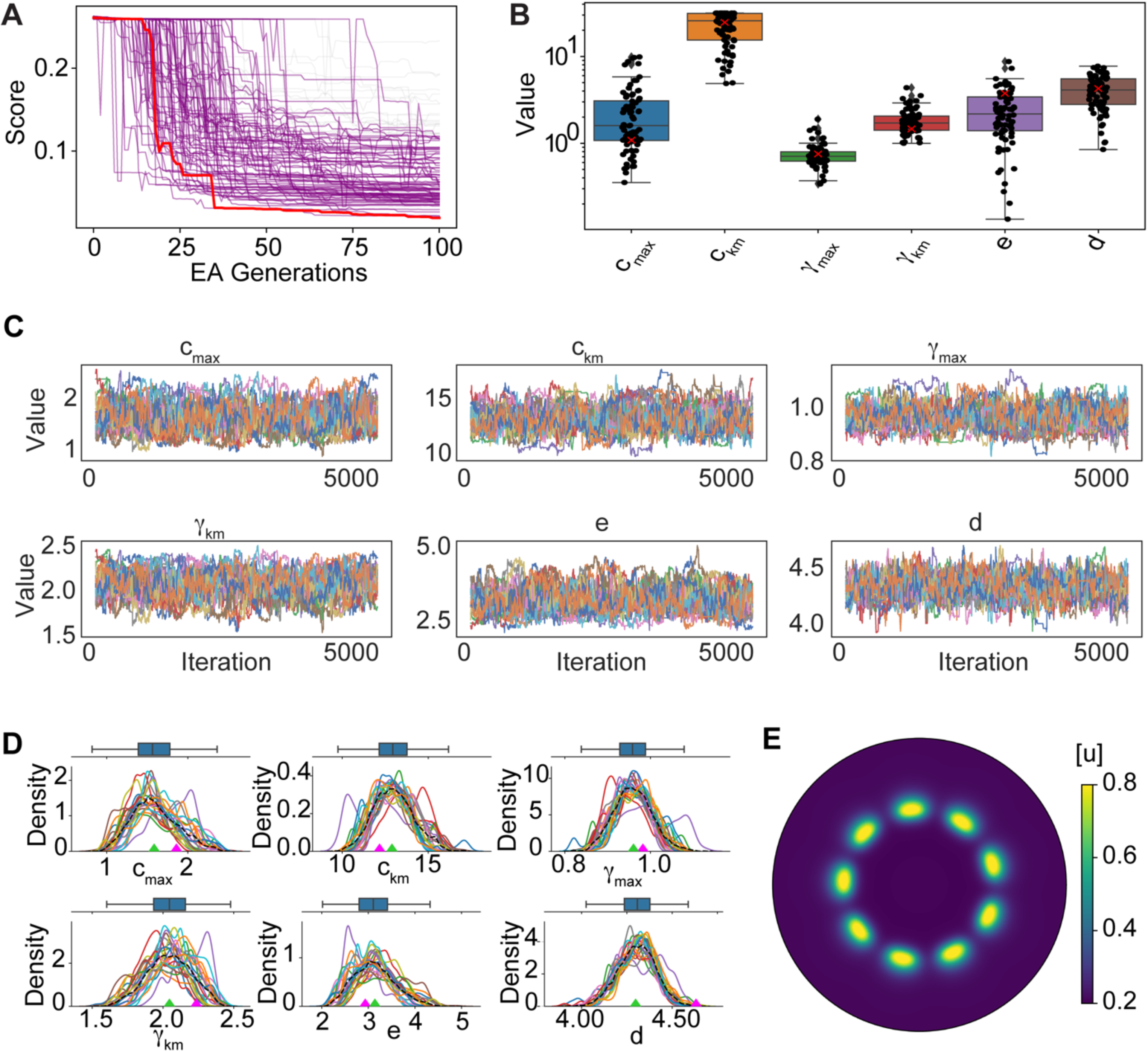
EA and DRAM-MCMC parameterization. **A)** EA parametrization runs (99 total). Best individual run shown in red and the runs that resulted in GTPase rosettes shown in purple. **B)** Individual parameter distributions from the successful EA runs shown in **A**. The best performing parameter set shown by red crosses. **C)** DRAM-MCMC chains for individual parameters post burn-in phase. **D)** Individual parameter densities for the chains shown in **C**. Representative parameter set values shown by green diamonds (Table 2). The worst scoring parameter set shown by magenta diamonds (Table S1). **E)** Active GTPase concentration for the worst scoring parameter set (Table S1).

**Supplemental Figure 4.**
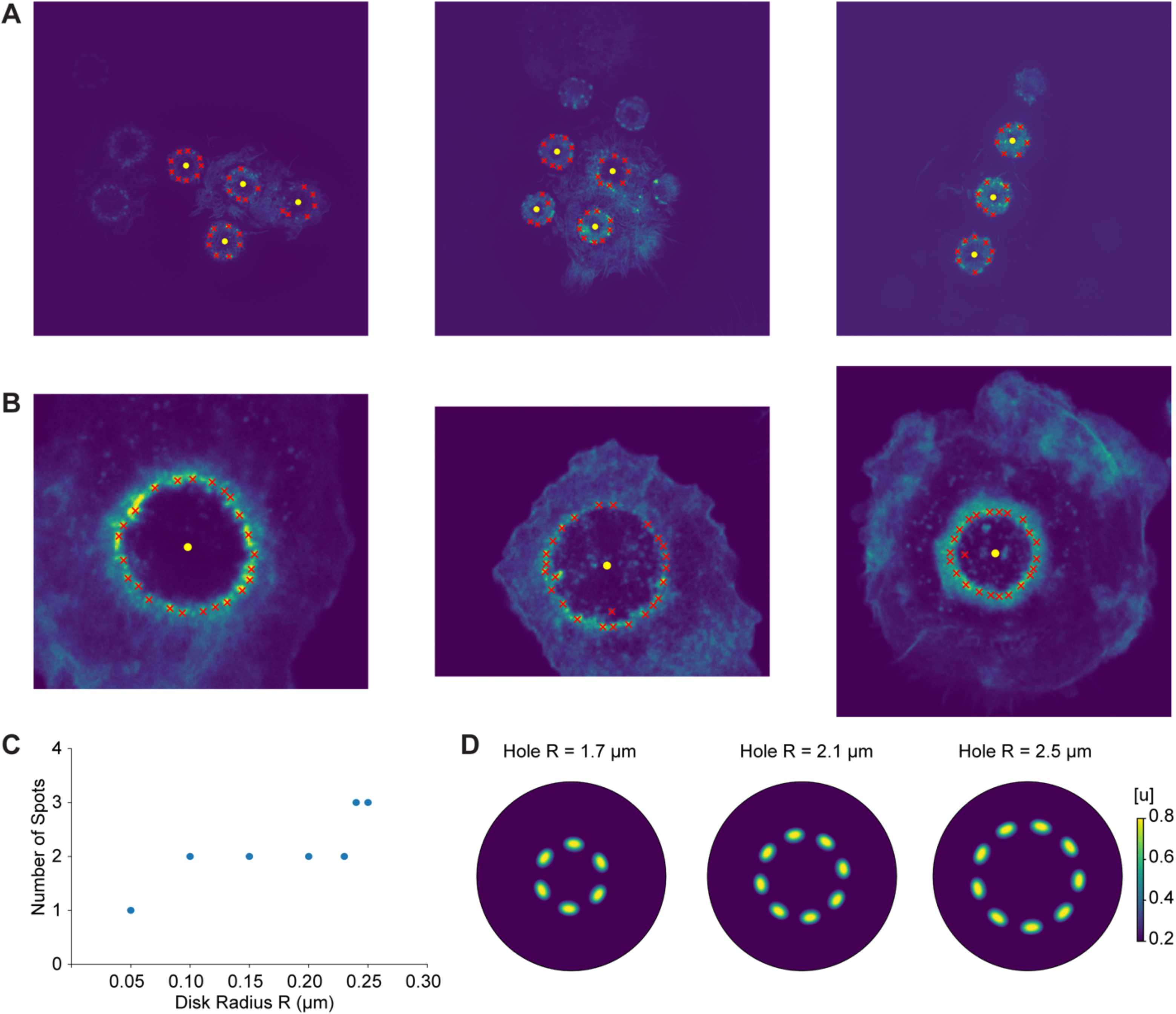
Additional information for experimental and simulated results on varying disk and hole sizes. **A)** Representative experimental results for disks of radius 1.75 μm. Podosomes are indicated with red circles. **B)** Same as A but using disks of radius 5 μm. **C)** Number of GTPases spots versus disk radius for small disks with radius less than 0.25 μm. **D)** Simulations for the representative parameter set (Table 2, negative *c_km_* and *γ_km_*) when changing the hole size.

**Table S1.**
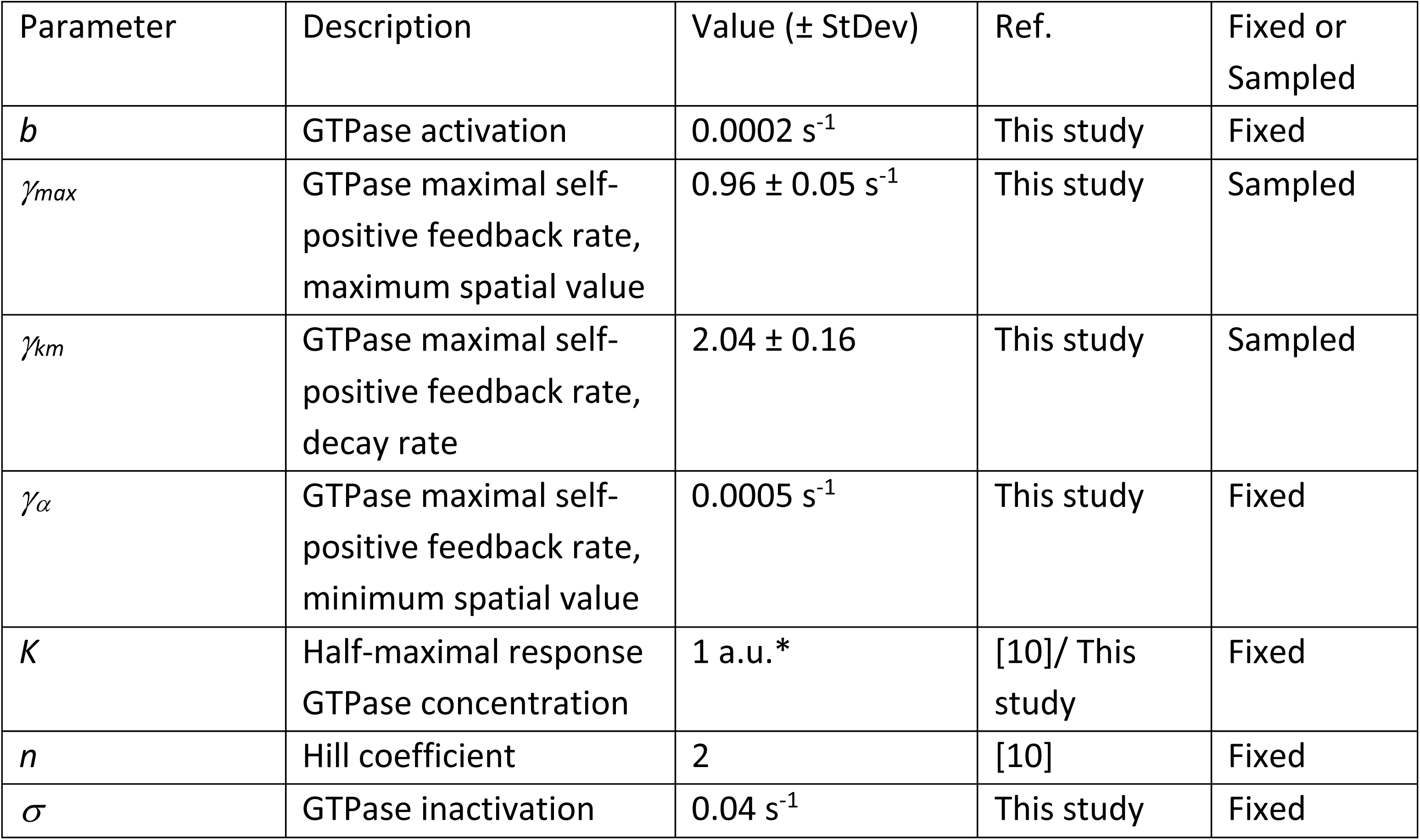
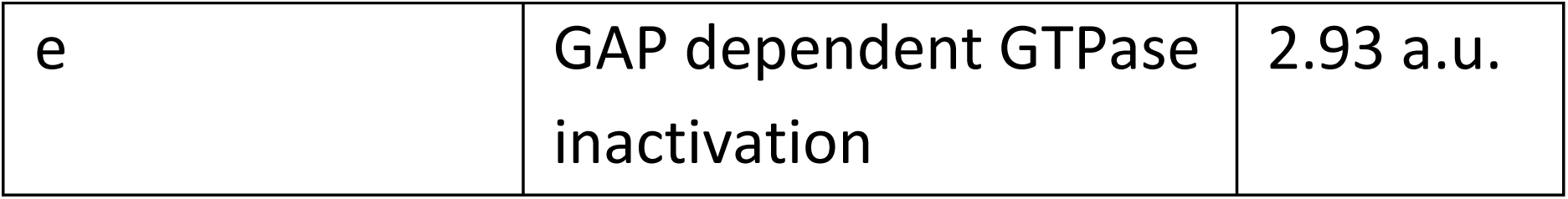
Worst scoring parameter set after running the MCMC. Fixed values same as in Table S2.

